# Targeting *ZC3H12C* improves T cell persistence and antitumor function in adoptive T cell therapy

**DOI:** 10.64898/2026.04.30.721618

**Authors:** Gayatri Kavishwar, Markus Perl, Christoph Heuser-Loy, Leonard Knoedler, Dhyani Shah, Fabio Mastrogiovanni, Konstantin Herfeld, Pedro Noronha, Manuela Kovács-Sautter, Maria Krieger, Stefan Loipfinger, Caio Raony Farina Silveira, Sascha Göttert, Roland Schelker, Luisa-Marie Becker, Timea Vadász, Carmen Gerlach, Gloria Lutzny-Geier, Luca Gattinoni, Hendrik Poeck, Christian Schmidl

## Abstract

Adoptive T cell therapy (ACT) has achieved remarkable clinical responses in hematologic malignancies but remains limited by progressive T cell dysfunction under chronic antigen stimulation. Here, we identify *ZC3H12C* as a conserved feature of dysfunctional T cells and show that its disruption enhances the durability and antitumor activity of engineered T cells. By integrating single-cell chromatin accessibility and transcriptomic profiling of human tumor-infiltrating lymphocytes (TILs), we identified the *ZC3H12C* locus as selectively remodeled in exhausted T cells. *ZC3H12C* induction is largely absent across acute T cell activation contexts, indicating regulation that is specific to chronic antigen-driven dysfunction. Genetic disruption of *ZC3H12C* improves T cell expansion, cytotoxicity, and expression of effector molecules during repeated *in vitro* stimulation, translating into enhanced tumor control *in vivo* across both T cell receptor (TCR) and chimeric antigen receptor (CAR) T cell therapy platforms. Improved efficacy is observed in hematologic, solid, and metastatic tumor models and is accompanied by increased T cell persistence. Further, *ZC3H12C* is enriched in clinical pre-infusion CAR T cell products associated with non-response. Together, these findings identify *ZC3H12C* as a T cell dysfunction-specific target to improve ACT performance.

## Introduction

ACT has emerged as a promising therapeutic intervention in cancer, achieving durable clinical responses particularly in hematologic malignancies^1,2^. CAR T cell therapies, for example, have demonstrated efficacy in otherwise refractory diseases, including chronic lymphocytic leukemia, B cell lymphomas, and multiple myeloma, inducing partial to complete remissions in a substantial fraction of patients^3–5^. Nonetheless, these successes remain limited: many patients fail to respond, and among initial responders, relapse is common. This challenge is especially pronounced in settings characterized by persistent antigen exposure - most notably in solid tumors, which account for the majority of human cancers, but also in subsets of hematologic malignancies^6^. Although cancer-directed CAR T cells and transgenic TCR T cells recognize tumors, clinical benefit is often transient and durable responses under sustained antigen stimulation remain rare^1,6^.

The limited efficacy of ACT across disease contexts is closely associated with the acquisition of dysfunctional T cell states, often referred to as T cell exhaustion^7^. Chronic antigen exposure, frequently accompanied by immunosuppressive cues in the tumor microenvironment, drives the progressive emergence of dysfunctional T cell states characterized by reduced cytokine production, impaired proliferative capacity, sustained expression of inhibitory receptors, and diminished cytotoxic function^8,9^. Such states correlate with poor tumor control and are commonly observed TILs and in engineered T cell products following persistent antigen stimulation^6,10–12^. While multiple transcriptional and regulatory features have been associated with exhausted T cells^7^, the molecular determinants that limit T cell fitness under chronic antigen exposure remain incompletely defined, and targeting these to improve ACT efficacy remains a challenge.

Single-cell RNA sequencing (scRNA-seq) has been widely used to characterize transcriptional programs associated with functional and dysfunctional T cell states^13–16^. However, terminal exhaustion is not defined at the level of gene expression alone, but is accompanied by durable epigenetic and gene-regulatory changes that are not readily inferred from RNA profiles^17^. Specifically, alterations in chromatin accessibility, histone modifications, and DNA methylation have been implicated in reinforcing dysfunctional T cell states that arise under chronic antigen exposure^18–21^. Consistent with this notion, immune checkpoint blockade can transiently induce effector-like transcriptional responses in exhausted T cells without substantially altering the underlying chromatin landscape, resulting in incomplete and temporally limited functional reinvigoration^22^. Genome-wide omics studies further suggest that terminally exhausted CD8⁺ T cells across chronic infection and human cancers adopt conserved regulatory architectures that distinguish them from effector and memory populations across chronic infection and human cancers^10,23^. Taken together, these observations argue that genes coupled to exhaustion-specific epigenetic remodeling, rather than those identified solely by differential expression, are more likely to reflect deep molecular commitment to dysfunction. Integrative single-cell profiling of chromatin accessibility together with transcription therefore offers a principled strategy to prioritize candidate regulators embedded within state-specific enhancer programs^24^.

An obstacle in identifying actionable regulators of T cell dysfunction is that many genes and surface markers associated with exhaustion are also expressed in activated T cells upon acute antigen exposure. For example, PD-1 is rapidly upregulated following T cell priming and vaccination, where it functions as an activation-induced checkpoint that tunes the magnitude and duration of early immune responses, with sustained expression emerging only under persistent antigen exposure^25^. Accordingly, reliance on individual markers or candidate regulators derived from chronic settings alone may be insufficient to distinguish acute activation from chronic dysfunction, complicating identification of suitable targets for T cell engineering. This highlights the need to systematically contrast chronic dysfunction programs against diverse acute activation contexts to pinpoint targets that interfere with dysfunction, but not T cell activation^15^.

Identifying general regulators of T cell dysfunction is further complicated by the fact that the effects of perturbing exhaustion-associated pathways are highly context dependent, differing across experimental systems, antigen-targeting strategies, and therapeutic platforms. As an example, an *in vivo* CRISPR-Cas9 screen in BCMA CAR T cells for multiple myeloma revealed that regulators enhancing T cell expansion *in vitro* failed to confer fitness advantages *in vivo*, whereas distinct factors selectively constrained persistence and antitumor activity only in *in vivo* conditions^26^. These observations highlight an unmet need to identify cell-intrinsic regulators that are selectively associated with chronic antigen-driven dysfunction, are conserved across chronic stimulation contexts and therapy products, and enhance T cell fitness *in vivo* without impairing essential activation programs. Such regulators would be particularly relevant in settings of chronic antigen exposure, including but not limited to solid tumors, where engineered T cells must retain proliferation capacity, effector function, and persistence under prolonged antigen exposure for sustained clinical benefits.

Therefore, to identify exhaustion-selective regulators that are cell-intrinsic, non-overlapping with activation, and context-agnostic, we focused on genes coupled to exhaustion-specific enhancer remodeling in human TILs, as opposed to evaluating expression changes alone. To this end, we applied our recent integrated single-cell chromatin accessibility and scRNA-seq dataset of human CD8⁺ TILs across multiple cancer entities^24^, to prioritize loci showing coordinated chromatin remodeling and transcriptional induction in terminally dysfunctional T cells. Using this approach, we identified *ZC3H12C* (encoding for Regnase-3) as a major remodeled locus in exhausted T cells. By comparing our dataset to T cells from *in vitro* activation and acute infection, we demonstrate that *ZC3H12C* is a marker specifically associated with T cell exhaustion, rather than general activation. Consistently, *ZC3H12C* was induced in T cells mounting chronic but not acute immune responses across a variety of murine models, supporting that programming of the *ZC3H12C* locus is exhaustion-specific. Further, we could show that genetic ablation of *ZC3H12C* enhanced proliferation, survival, and effector function of engineered T cells *in vitro*, which translated into improved *in vivo* tumor control in several adoptive therapy models. Taken together, we identified *ZC3H12C* as a general regulator of antitumor T cell function specific to chronic antigen-exposure settings, which therefore represents a potent cross-platform target to improve T cell-based immunotherapies.

## Results

### Human TIL chromatin accessibility and gene expression profiles uncover exhaustion-specific reprogramming of the *ZC3H12C* locus

To identify broadly acting, stably reprogrammed regulators of T cell dysfunction that minimally overlap with activation, we first analyzed our integrated single-cell chromatin accessibility and transcriptome dataset of human CD8⁺ TILs from hepatocellular carcinoma, renal cell carcinoma, head and neck squamous cell carcinoma, and basal cell carcinoma^24^. Across all tumor types, we detected dysfunctional TIL states and a shared set of genes with distinct chromatin remodeling relative to functional TILs. Joint analysis of chromatin and scRNA-seq profiles enabled inference of cell state-specific enhancer regulation of key TIL genes. We reasoned that a ranking of genes based on such integration of chromatin and expression data highlights candidates that undergo deep molecular rewiring. Therefore, we revisited the integrated multi-omics-based ranking of genes in terminally exhausted T cells (T_EX-term_, modular nomenclature^27^: TX_t_+), where we found *ZC3H12C* as a top-ranked candidate in solid cancers in addition to known genes such as *TOX* and *ENTPD1* (**Figure 1A**). Indeed, *ZC3H12C* had similar high chromatin accessibility-based gene activity in T_EX_ clusters 1 and 2 as observed for *TOX*, *ENTPD1* (encoding CD39), *CTLA4*, and *HAVCR2* (encoding TIM3), separating them well from T cell precursor exhausted cells (T_PEX_, modular nomenclature: TX_p_+) (high *TCF7* activity, cluster 3), functional, minimally differentiated memory T cells (high *BACH2* activity, no *TOX*, cluster 4), and functional, highly differentiated cytotoxic effector T cells (high *CX3CR1* activity, no *TOX*, cluster 6) (**Figure 1B-C**). In agreement with increasing chromatin accessibility, we observed elevated *ZC3H12C* mRNA levels in T_EX_ clusters (**Figure 1D**). Visual inspection of the *ZC3H12C* locus showed a profound remodeling of the surrounding chromatin landscape with a large fraction of accessible chromatin sites that are predicted to act as enhancers regulating *ZC3H12C* expression (**Figure 1E**, left panel). Interestingly, the chromatin remodeling at the *ZC3H12C* locus is highly specific for T_EX-term_ (clusters 1 and 2), similar to what is observed for loci such as *ENTPD1* (**Figure 1E**, right panel) and *TOX* (**Figure S1A**).

**Figure 1:**
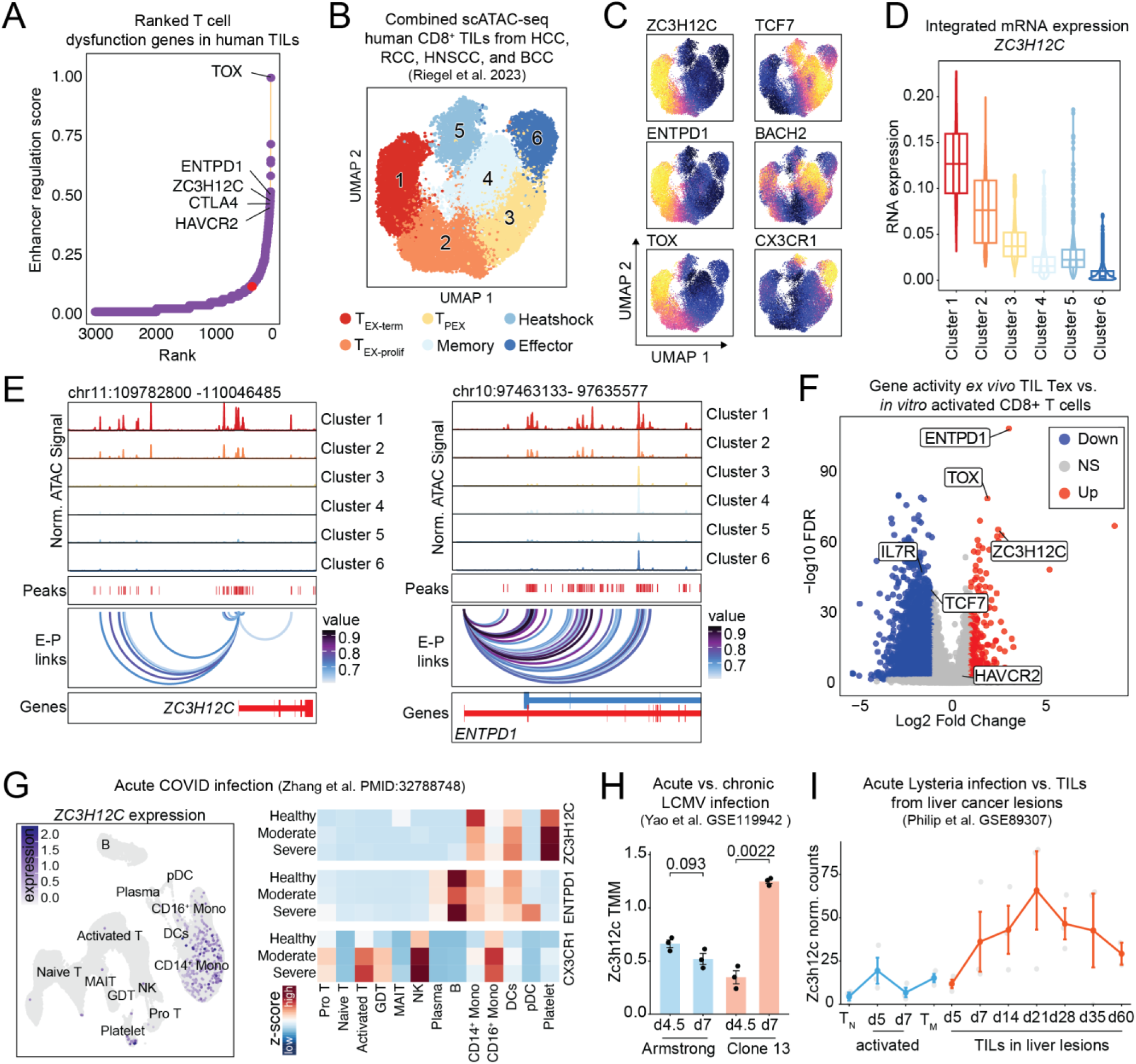
*ZC3H12C* is specifically induced in dysfunctional T cells. **(A)** Enhancer regulation score ranking of T cell dysfunction-associated genes in T_EX-term_ based on combined single-cell chromatin accessibility and transcriptome profiles from human TILs from Riegel et al.^24^. **(B)** UMAP projection with clustering and **(C)** chromatin-accessibility-based gene activity for specific marker genes in TILs. **(D)** mRNA expression of *ZC3H12C* in clusters of (B). **(E)** Genome browser track visualization containing normalized chromatin accessibility (scATAC-seq) summarized for each cluster of (B), accessible regions (’peaks’), predicted regulation of gene promoters by distal enhancers (‘E-P links’) with a value score indicating the correlation between accessibility of an enhancer and the expression of the corresponding gene, and the gene structure (’genes’) for the *ZC3H12C* and *ENTPD1* loci. **(F)** Comparison of chromatin-accessibility-based gene activity of T_EX-term_ from tumors against *in vitro* activated CD8^+^ T cells from healthy donors that were treated with TransAct™ in the presence of IL-2 for 18 hours to 3 days. **(G)** *ZC3H12C* expression in PBMCs during acute COVID-19 infection (left panel), and z-score normalized expression together with *ENTPD1* and *CX3CR1* for each cell population in the blood, grouped in healthy donors, moderate, and severe COVID-19 disease from Zhang et al.^28^. **(H**) *Zc3h12c* expression in virus-specific T cells at day 4.5 and day 7 during LCMV infection from Yao et al.^30^ **(I)** *Zc3h12c* expression of antigen-specific naïve (T_N_), effector, and memory T (T_M_) cells in a model of acute Listeria infection, as well as *Zc3h12c* expression in chronically activated tumor-specific T cells isolated from lesions of an autochthonous liver cancer model from Philipp et al.^10^. Statistical significance was assessed using a two-tailed unpaired Student’s t-test for pairwise comparisons. Normalized *Zc3h12c* expression is shown as mean ± SEM.

As T cell activation and dysfunction programs are to some degree overlapping, we next compared chromatin-based gene activities between dysfunctional CD8^+^ TILs and short-term *in vitro* activated human CD8^+^ T cells from peripheral blood mononuclear cells (PBMCs) isolated from healthy donors. Interestingly, we observed significant higher activity of *ZC3H12C* along with *TOX* and *ENTPD1* in dysfunctional cells compared to *in vitro* activated T cells, while *HAVCR2*, which is associated with both activation and dysfunction, was not significantly different in this comparison (**Figure 1F**). Analysis of a dataset of acute COVID-19 infection^28^ confirmed that similar to *ENTPD1*, *ZC3H12C* was not expressed in activated T cells of any disease severity, but was expressed in myeloid cells as reported previously^29^ (**Figure 1G**, **Figure S1B**). However, markers of effector function including *CX3CR1*, *PRF1*, *GZMB*, *IFNG*, and activation, including *PDCD1* and *HAVCR2*, were upregulated in T cells from COVID-19 patients (**Figure 1G**, **Figure S1C**), indicating that acute immune responses do not induce significant levels of *ZC3H12C* mRNA in human T cells. We observed similar specificity of *ZC3H12C* expression also in murine models of infection and cancer. While *Zc3h12c* was not induced upon Lymphocytic Choriomeningitis Virus (LCMV) infection with the Armstrong strain, which causes acute infection, *Zc3h12c* was significantly induced on day 7 post infection with LCMV clone 13, which causes chronic infection^30^ (**Figure 1H**). Further, *Zc3h12c* was also not induced during acute Listeria infection in effector and memory T cells, while it was upregulated in TILs isolated from an autochthonous liver cancer with a peak of expression three weeks after tumor onset^10^ (**Figure 1I**). In summary, we observed profound chromatin remodeling of the *ZC3H12C* locus with increased expression in TILs when they developed towards a dysfunctional state, and this induction is specific to T cells in chronic activation settings but not observed in acute T cell responses.

### Deletion of *ZC3H12C* improves expansion, cytotoxicity, and effector molecule expression in TCR-engineered T cells

We reasoned that due to its specific remodeling in dysfunctional T cell states, *ZC3H12C* may be a potent target to improve T cell immunotherapy products. We therefore used CRISPR gene editing to delete *ZC3H12C* in primary human CD8^+^ T cells that were transduced with the NY-ESO-1-specific transgenic TCR^31^, which recognizes an intracellular cancer-testis antigen via peptide-HLA recognition. After knockout and transduction, cells were expanded in the presence of IL-2, with media changes every second day for up to 25 days. We observed significantly higher cell numbers in the TCR T cell product harboring the *ZC3H12C* knockout compared to the CRISPR control that is targeting the *AAVS1* region, a safe harbor locus in the genome (**Figure 2A**). Next, we repetitively co-cultured NY-ESO-1 TCR T cells with antigen-expressing target cells (SK-MEL-23, a cutaneous melanoma cell line) for 48h to induce T cell mediated cytotoxicity. The co-culture was subsequently repeated for up to five rounds to cause chronic TCR activation. TCR T cell and target tumor cell numbers were adjusted to the same effector to target ratio of 1:1 after each round of co-culture. Again, we observed a higher cell count in the *ZC3H12C* knockout TCR T cells than in the CRISPR control (**Figure 2B**). In addition to superior expansion, *ZC3H12C* knockout TCR T cells showed improved tumor cell lysis: using live-cell imaging, we observed that while CRISPR control cells lost their cytotoxic function, indicative of their progression to a dysfunctional state, the *ZC3H12C* KO TCR T cells maintained their ability of sustained target cell killing over several rounds of co-culture. Although the exact co-culture round when the control knockout cells lost their killing capacity was donor-dependent, *ZC3H12C* KO TCR T cells consistently maintained their cytotoxic capacity in ten biological replicates (**Figure 2C-E**, **Figure S2A-B**). Further, *ZC3H12C* knockout TCR T cells analyzed after repeated co-culture, following short term stimulation in the presence of protein transport inhibitors, showed significantly elevated levels of Interferon gamma (IFNγ), Granzyme B (GzmB), and Interleukin-2 (IL-2), and increased protein expression of the transcription factor T-bet (**Figure 2F, Figure S2C**). Interestingly, we did not observe a change in CD39 or TOX after three rounds of restimulation when cells of the control TCR population were still sufficient in numbers for analysis. Further, we observed a higher fraction of cells that co-expressed the two effector molecules IFNγ and GzmB, indicating higher polyfunctionality of engineered TCR T cells (**Figure 2G-H**). In summary, *ZC3H12C* knockout improved expansion, cytotoxicity, and expression of effector molecules in the TCR T cell product.

**Figure 2:**
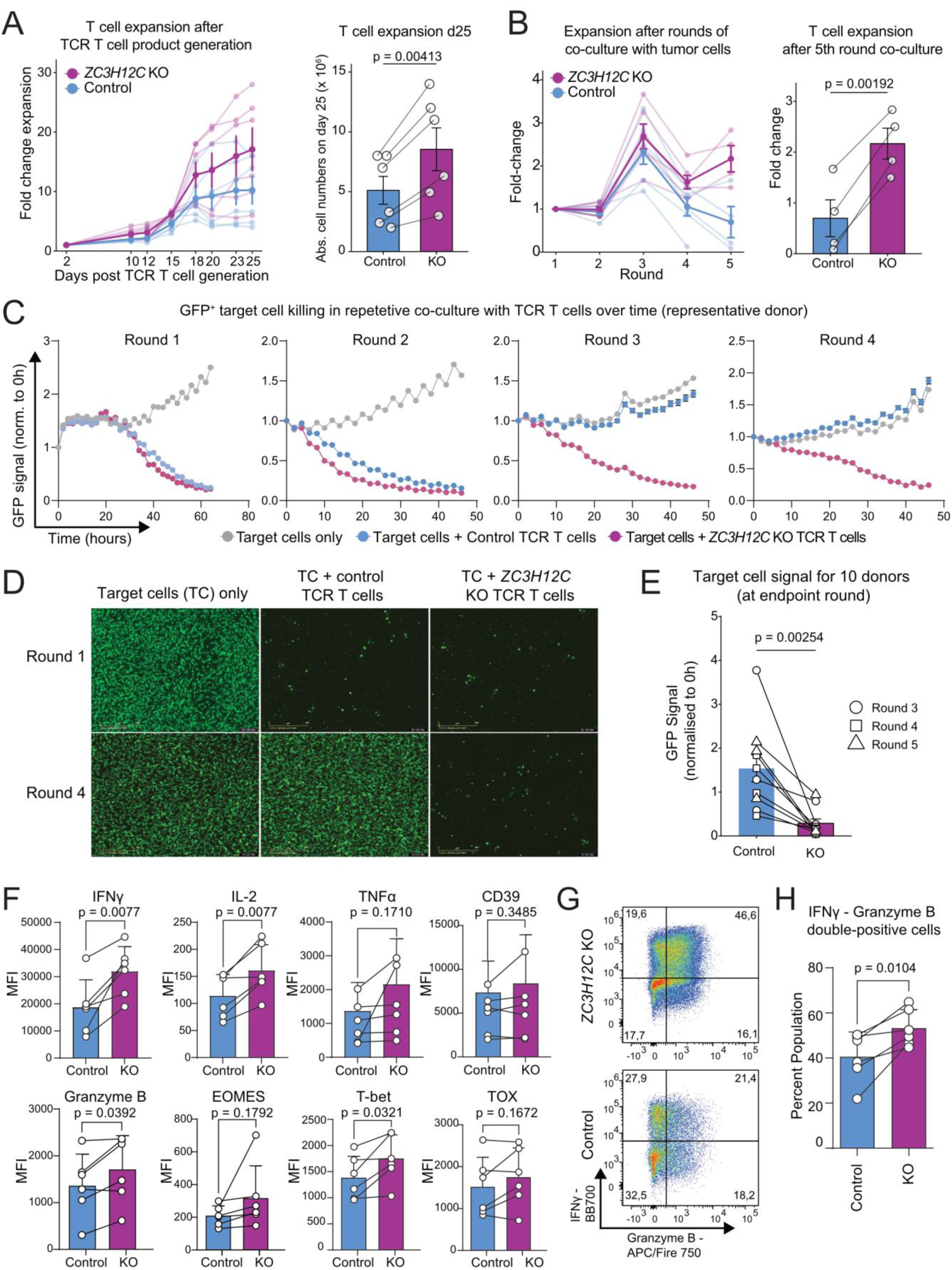
Deletion of *ZC3H12C* improves expansion, cytotoxicity, and effector molecule expression in TCR T cells. **(A)** Expansion of NY-ESO-1 TCR^+^ T cells with *ZC3H12C* KO (KO) or *AAVS1* targeting (Control). Fold-changes in respect to seeding cell numbers for up to 25 days of expansion are shown. **(B)** Expansion of NY-ESO-1 TCR^+^ T cell during a serial co-culture assay with SK-MEL-23 melanoma target tumor cells (E:T=1:1) for five consecutive rounds of antigen re-challenge. Co-cultures were maintained for 48 hours per round and monitored by live-cell imaging. After each round of co-culture, T cells were harvested, counted, and seeded onto freshly plated GFP^+^ SK-MEL-23 target cells, maintaining the E:T ratio of 1:1, to initiate the next round of antigen re-challenge. **(C)** Real-time cytotoxicity kinetics over four consecutive rounds of NY-ESO-1 TCR T cell co-culture with SK-MEL-23 melanoma target tumor cells (E:T=1:1). T cell-mediated cytotoxicity was quantified by loss of GFP signal from GFP^+^ SK-MEL-23 target cells. *ZC3H12C* KO cells are compared to Control *AAVS1* TCR T cells, tumor target cells only are shown as a further control. **(D)** Representative Incucyte images from the cytotoxicity assay in (C) showing GFP^+^ target cells at Round 1 and Round 4 of co-culture **(E)** Endpoint quantification of tumor cell killing after three, four, or five rounds of co-culture (*n* = 10 healthy donors). Endpoints were considered reached when *AAVS1* KO T cell numbers were insufficient to commence an additional round of co-culture. **(F)** Flow cytometry analysis of NY-ESO-1 TCR^+^ T cells collected following Round 3 of killing/re-stimulation. **(G)** Representative flow cytometry dot plots showing IFNγ and GzmB co-expression. **(H)** Summary of IFNγ and GzmB double-positive cells for all experiments. Statistical significance was determined using a two-tailed paired Student’s t-test. Bar graphs show mean ± SEM with individual data points representing independent donors.

### *ZC3H12C*-deleted NY-ESO1 TCR T cells show improved *in vivo* efficacy in a solid xenograft tumor model

Encouraged by the improved function of *ZC3H12C*-edited TCR T cells *in vitro*, we next tested their performance in a SK-MEL-23 melanoma xenograft model (**Figure 3A**). Here, 1 x 10^6^ NY-ESO-1^+^ SK-MEL-23 melanoma cells were subcutaneously transplanted into IL-15 producing NXG mice. 27 days later 2 x 10^6^ TCR T cells were adoptively transferred, and we monitored tumor growth, survival, and isolated peripheral blood to detect TCR T cells during treatment using flow cytometry. To avoid graft-versus-host reactions, we deleted the endogenous TCR in transferred T cells (**Figure 3B, Figure S3A**). In the final product, we achieved high CRISPR editing efficiencies, and we further purified cells expressing the NY-ESO-1 TCR by sorting before T cell transfer (**Figure 3C, Figure S3B-C**). Tumors grew most aggressively in mice receiving no T cells, with rapid and uninterrupted tumor outgrowth. Introduction of NY-ESO-1 TCR T cells with the *AAVS1* knockout improved tumor control relative to this baseline, indicating partial antitumor activity (**Figure 3D-E**). However, the most robust therapeutic effect was achieved by CRISPR knockout of *ZC3H12C* in NY-ESO-1 TCR T cells, with tumors showing the deepest and most consistent regression and a greater delay in progression across all mice. These observations were mirrored in significantly prolonged survival of *ZC3H12C* TCR T cell-treated mice in comparison to *AAVS1* and UT control groups (**Figure 3F**). As low persistence and expansion are limiting factors of cellular T cell therapies, we assessed the absolute numbers of NY-ESO-1 TCR T cells in the peripheral blood of mice after adoptive TCR T cell transfer. The *ZC3H12C*-ablated cells showed significantly higher numbers than *AAVS1* control TCR T cells at day 18 post-transfer, which coincided with the time window of maximal tumor reduction in the *in vivo* model (**Figure 3G-H, Figure S3D**). Together, in line with our *in vitro* findings, our *in vivo* results show that *ZC3H12C* deletion improves TCR T cell efficacy in a xenograft tumor model, probably driven by increased expansion and killing capacity of the engineered TCR T cell product.

**Figure 3:**
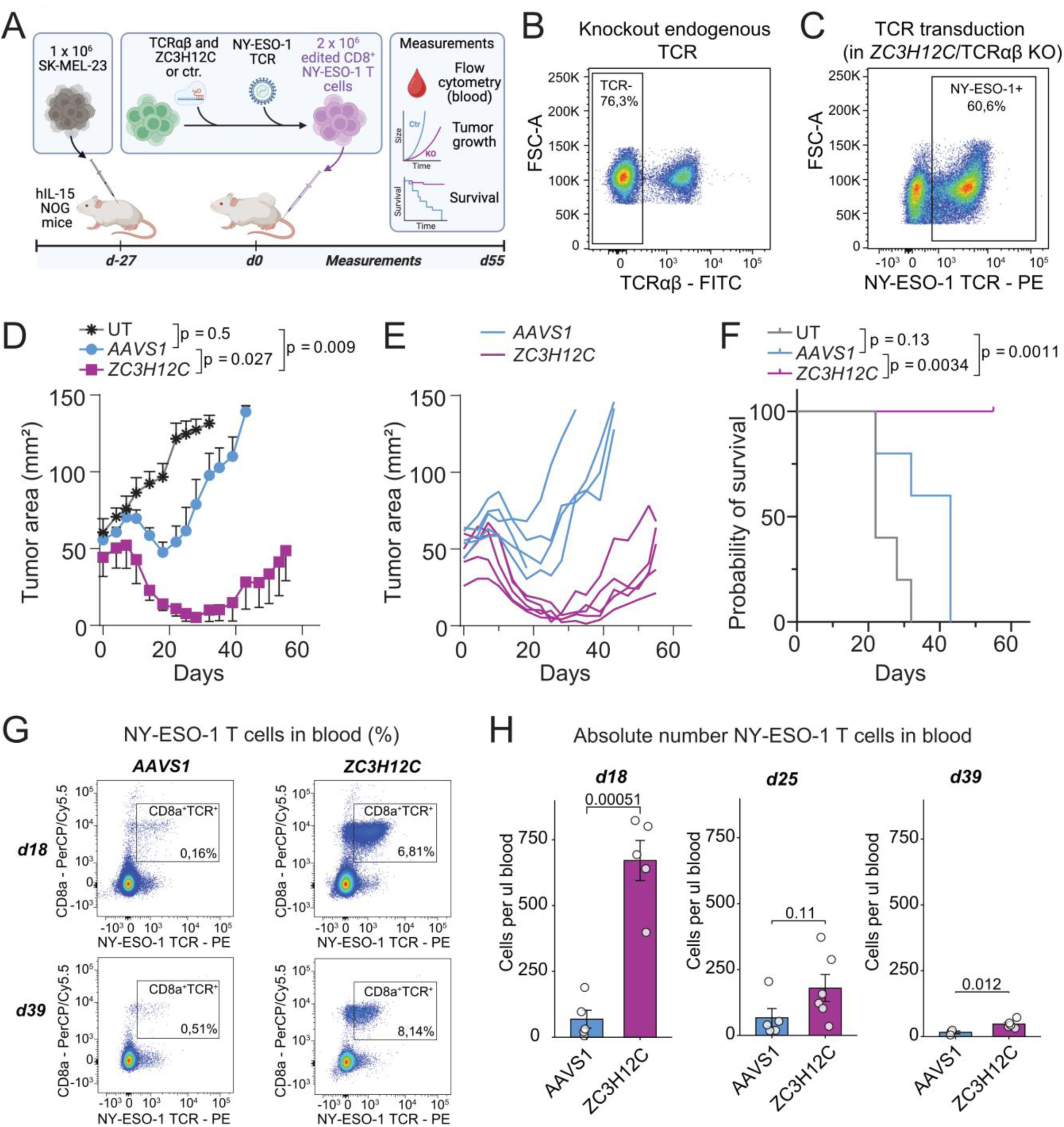
*ZC3H12C*-deleted TCR T cells show improved *in vivo* efficacy in a solid xenograft tumor model. **(A)** Schematic overview of the mouse model and treatment schedule used to evaluate NY-ESO-1 TCR T cell efficacy *in vivo*. **(B)** Editing efficiency of endogenous TCR knockout before NY-ESO-1 TCR T cell transduction. **(C)** NY-ESO-1 TCR transduction efficiency in engineered T cells prior to adoptive transfer. **(D)** Cumulative tumor growth kinetics following treatment with NY-ESO-1 TCR T cells, comparing *ZC3H12C* KO versus *AAVS1* targeting control and untreated mice (UT). **(E)** Individual tumor growth curves for each mouse from (D). **(F)** Kaplan-Meier survival analysis of mice from the tumor experiment in (D). **(G)** Representative flow cytometry dot plots illustrating the percentage of circulating NY-ESO-1 TCR^+^ T cells in peripheral blood at day 18 and day 39 post-adoptive TCR T cell transfer. **(H)** Absolute cell numbers of NY-ESO-1 TCR T cells in peripheral blood at day 18, day 25 and day 39 post-adoptive TCR T cell transfer. Statistical significance was assessed using a two-tailed unpaired Student’s *t*-test for pairwise comparisons. Peripheral blood T cell counts are shown as mean ± SEM. Survival was analyzed using the log-rank (Mantel-Cox) test.

### *ZC3H12C* knockout improves survival in a hematologic and an experimental metastatic CAR T cell tumor model

Having shown improved function of *ZC3H12C*-deleted therapeutic TCR T cells *in vitro* and *in vivo*, we next tested the broader applicability of *ZC3H12C*-targeting in additional models of adoptive T cell therapies using CAR T cells. First, we employed an aggressive liquid tumor model. Specifically, 0.5 x 10^6^ NALM6 B-cell precursor acute lymphoblastic leukemia cells were injected in NXG mice (**Figure 4A**). Three days after tumor engraftment, 1 x 10^6^ CD19 CAR T cells with a CD28 co-stimulatory domain, with either *ZC3H12C* deletion or CRISPR control, were transferred into leukemia-bearing mice. In this model, *ZC3H12C* KO CD19 CAR T cells provided a significant survival benefit and lowered the tumor burden in comparison to CRISPR control CD19 CAR T cells (**Figure 4 B-C**). To determine whether the benefits of *ZC3H12C*-deletion extend beyond hematologic malignancies in CAR T cell models, we next wanted to evaluate its impact in a non-hematopoietic, solid tumor-like context. To this end, we used HER2 CAR T cells with a 4-1BB costimulatory domain in an experimental lung metastasis model (**Figure 4D**). In this system, 1 x 10^6^ SKOV3 ovarian adenocarcinoma cells were injected intravenously, leading to efficient lung colonization and progressive pulmonary tumor growth. Seven days after tumor injection, when the pulmonary metastases were established, as confirmed by bioluminescence measurement, 0.1 x 10^6^ HER2 CAR T cells were adoptively transferred. Similar to the NALM6 model, animals that were treated with *ZC3H12C* KO HER2 CAR T cells had a significant survival benefit and delayed tumor progression compared to mice treated with CRISPR control HER2 CAR T cells (**Figure 4E-F**). Collectively, the improved performance of *ZC3H12C*-deficient CAR T cells in both hematologic and non–hematopoietic metastasis models indicates that *ZC3H12C* targeting can broadly augment ACT, including in more challenging solid tumor-like contexts.

**Figure 4:**
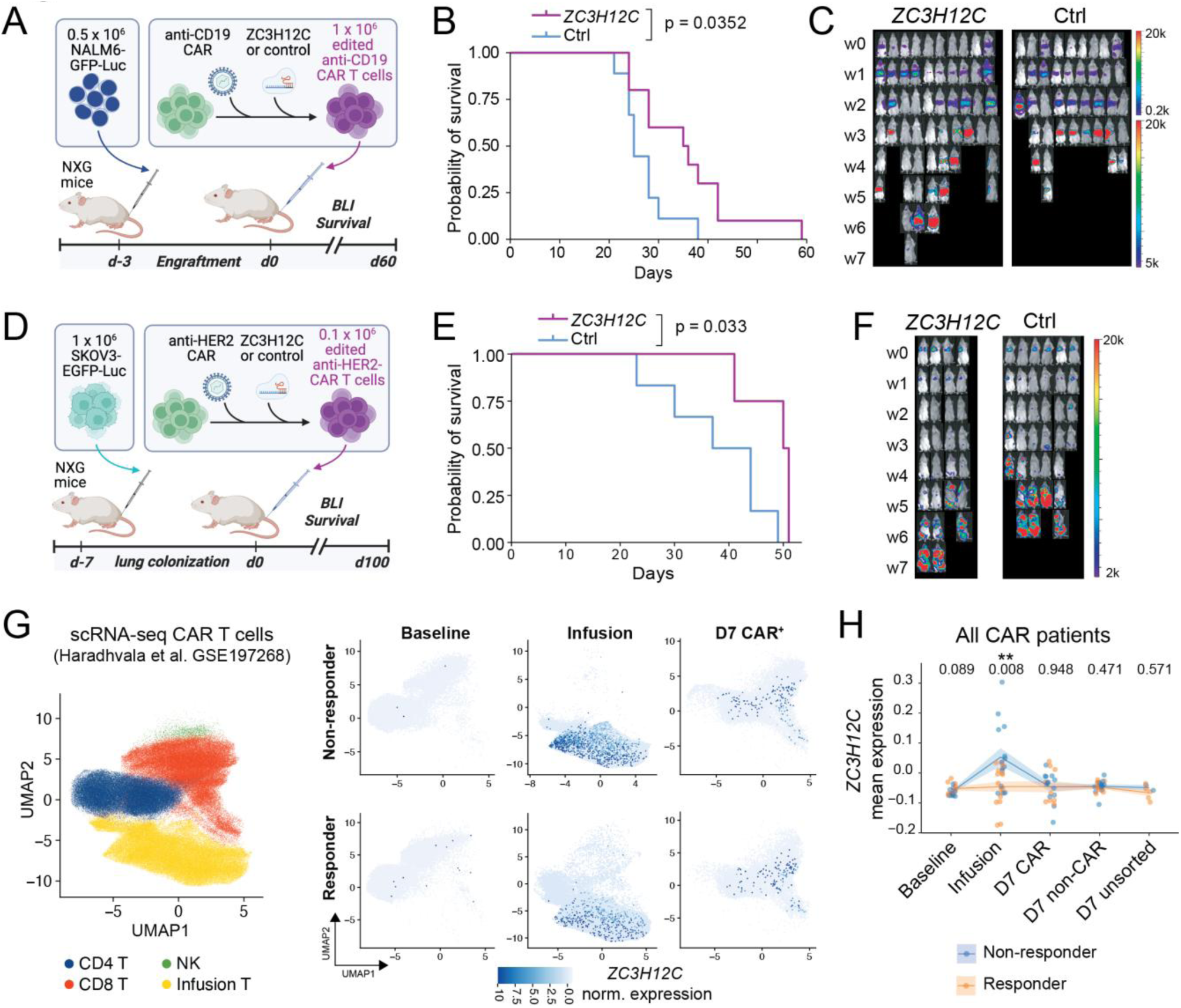
*ZC3H12C* knockout improves survival in two CAR T cell tumor models and is upregulated in pre-infusion CAR T cell products of non-responding patients. **(A)** Schematic overview of the acute lymphoblastic leukemia xenograft model used to evaluate the *in vivo* efficacy of human *ZC3H12C*-deleted CD19 CAR T cells. **(B)** Kaplan-Meier survival analysis of mice treated with CD19 CAR T cells comparing *ZC3H12C* KO versus CRISPR control CD19 CAR T cells. **(C)** Bioluminescence images (BLI) of leukemia burden in mice from the CD19 CAR T cell model in (B). **(D)** Schematic overview of the human HER2 CAR T cell xenograft model. **(E)** Kaplan-Meier survival analysis of mice treated with HER2 CAR T cells comparing *ZC3H12C* KO HER2 CAR T cells with CRISPR control HER2 CAR T cells. **(F)** Representative IVIS bioluminescence images showing tumor burden in the HER2 CAR T cell model from (E). **(G)** UMAP projection of CAR T scRNA-seq data from Haradhvala et al.^32^, showing *ZC3H12C* expression stratified by clinical response (Responders versus Non-responders) in baseline (T cells after isolation), the CAR T cell infusion product (Infusion), and in CAR T cells isolated from patients seven days after infusion (D7 CAR). (H) *ZC3H12C* expression across all patients from (G). Each point represents the mean pseudobulk expression of a patient at each timepoint. Statistical significance was assessed using a two-sided unpaired Wilcoxon rank-sum test. Survival was analyzed using the log-rank (Mantel-Cox) test.

### *ZC3H12C* is upregulated in pre-infusion CAR T cell products of non-responding patients

As our data suggested that *ZC3H12C* negatively impacts adoptive cell therapy products, we investigated a possible correlation of *ZC3H12C* expression with adoptive T cell therapy outcomes. To this end, we re-analyzed a previously published scRNA-seq dataset of CAR T cells manufactured for cancer patient treatment, and from which expression data of the isolated donor T cells (“baseline”), infused CAR T cells (“infusion”), and re-isolated CAR T cells from blood (“d7 CAR”) were obtained, and information on response to therapy was available^32^. The dataset comprises 32 patients with large B cell lymphoma treated with either axicabtagene ciloleucel (axi-cel) or tisagenlecleucel (tisa-cel). *ZC3H12C* expression was not detectable in baseline donor T cells from leukapheresis, however, it was expressed in the infusion product that underwent activation during manufacturing (**Figure 4G**). Here, *ZC3H12C* expression was significantly higher in CAR T cells from non-responding patients in comparison to CAR T cells that yielded a clinical response (**Figure 4H, Figure S4A**). A similar pattern was observed for *CTLA4* expression, but not for other individual genes including *TOX*, *ENTPD1*, or *PDCD1* (**Figure S4B**). In summary, our analysis shows that *ZC3H12C* expression in CAR T cell products is negatively associated with clinical responses.

## Discussion

Adoptive T cell therapy has transformed the treatment of several hematologic malignancies, but its broader impact remains limited due to the progressive loss of T cell function under chronic activation due to persistent antigen exposure. Here, we identify *ZC3H12C* induction as a conserved and selective feature of dysfunctional T cells and demonstrate that genetic disruption of this locus enhances the durability and antitumor activity of engineered T cells *in vitro* and across multiple therapeutic platforms *in vivo*. Using cell-state-specific regulome data from integrated single-cell chromatin accessibility and transcriptomic profiling of human TILs, we identified *ZC3H12C* as a candidate gene that is specifically and profoundly remodeled in terminally exhausted TILs. Strikingly, *ZC3H12C* induction was largely absent across diverse acute activation and acute infection contexts, suggesting that its regulation is not a general feature of T cell activation but rather reflects chronic antigen-driven dysfunction. We demonstrate that *ZC3H12C*-ablation enhances T cell expansion and serial tumor challenge performance *in vitro*, which translated to improved tumor control in both TCR T and CAR T *in vivo* models, independent of the costimulatory domain used in the CAR. Notably, *ZC3H12C* targeting improved efficacy not only in a hematologic model but also in primary and metastatic solid tumor settings.

A key challenge in identifying actionable genes involved in T cell dysfunction is that transcriptional signatures of exhaustion overlap with those induced during acute activation^15,33^. For example, inhibitory receptors such as PD-1 or TIM3 are transiently induced by strong TCR signalling, which complicates their interpretation as exhaustion-specific markers^34,35^. Deleting genes involved in activation may disrupt physiological processes that are essential for antitumor efficacy as constitutive loss of PD-1 signalling can impair optimal CD8 memory formation, recall, and metabolic homeostasis required for long-lived quiescent memory^36–40^. In contrast, our enhancer-focused approach allowed prioritizing genes that not only increase in expression in dysfunctional T cells but also exhibit specific and profound chromatin accessibility changes of linked regulatory elements, a hallmark of cell state commitment. This strategy leverages the observation that exhaustion is accompanied by durable changes in chromatin accessibility and enhancer usage, which are incompletely captured by RNA profiles alone^10,17,20^. Consistent with prior work showing that exhausted T cells acquire a distinct chromatin or DNA methylation landscape that limits durable reinvigoration by checkpoint blockade^19,22^, the identification of *ZC3H12C* through exhaustion-specific enhancer regulation suggests that this locus may be part of the deeper regulatory architecture that constrains T cell function during chronic antigen exposure.

By systematically comparing tumor and infection settings with chronic T cell activation to acute activation conditions, we further validated the dysfunction-specificity of *ZC3H12C*. For example, in severe COVID-19 infection, T cells upregulated effector molecules including *GZMB*, terminal differentiation markers such as *CX3CR1*^41^, shared exhaustion/activation markers including *PDCD1* and *HAVCR2*, but did not upregulate *ZC3H12C*^28^. Further, in single-cell datasets from acute viral infection, *ZC3H12C* expression remained low, whereas robust induction was observed in chronic murine infection (LCMV Clone 13) and in CD8⁺ TILs in an autochthonous solid tumor model^10,30^. These cross-species patterns argue that *ZC3H12C* is regulated by conserved pathways engaged during persistent antigen exposure rather than by activation alone. Such selectivity is particularly relevant for ACT engineering, where interventions that blunt activation programs can impair initial expansion and antitumor responses^42,43^. Our findings therefore support the concept that filtering candidate genes against acute antigen activation contexts can enrich for targets that preferentially modulate chronic dysfunction without compromising essential activation responses. Interestingly, TILs with high *ZC3H12C* expression also express *ENTPD1* (CD39), which is a marker for tumor-reactive T cells^44^, further supporting chronic antigen stimulation in the clusters that we used to rank *ZC3H12C* as a candidate gene. To our knowledge, *ZC3H12C* has not been explored as a target for T cell engineering so far. Although our analysis confirms induction also in mouse models of T cell exhaustion in both infection and cancer in existing datasets, it may be that due to model system- or species-dependent differences in expression that in previous studies *ZC3H12C* was ranked lower than other candidates for downstream validations compared to our ranking, which is based on integrated multiomics data and was obtained from primary human samples.

In our work, disruption of *ZC3H12C* consistently enhanced T cell expansion *in vitro*, suggesting that its expression retains proliferation during chronic stimulation and may limit the accumulation of functional effectors. In addition to enhanced expansion, we observed a robust increase in cytotoxicity and effector molecule production (IFNγ and GzmB) in engineered T cells, consistent with improved functional competence rather than simply increased proliferation. Exhausted T cells can retain partial cytotoxic programs but often show impaired cytokine production and proliferative renewal, hence improving both axes is particularly desirable for durable tumor control^8^. The improved antitumor efficacy observed with engineered NY-ESO-1 TCR T cells demonstrates that targeting *ZC3H12C* can enhance antitumor immunity *in vivo*, supporting relevance to TCR-engineered therapies where chronic antigen exposure is prevalent^31^. The improved efficacy is at least in part driven by the improved expansion and persistence *in vivo*, as we observed significantly more TCR T cells in the peripheral blood at the time of strongest tumor regression. The consistent benefit in two independent CAR T models, comprising NALM6 leukemia and SKOV3 lung metastasis, suggests that *ZC3H12C* disruption may provide a broadly applicable enhancement strategy across CAR designs with different co-stimulatory domains as well as across tumor contexts.

Emerging evidence indicates that the cell states imposed by *ex vivo* activation and transduction protocols critically shape the fitness and exhaustion program of engineered T cells^45–47^. We observe increased expression of *ZC3H12C* in the pre-infusion product of CAR T cells, where it was expressed higher in CAR T cells for patients who did not respond to therapy. We speculate that prolonged T cell activation during the manufacturing process to achieve efficient transduction is therefore one factor that contributes to upregulating *ZC3H12C* expression. This may negatively affect the cell product performance even before infusion, potentially in addition to contributing to exhaustion *in vivo*. Although this observation needs further investigation, it may open the possibility to transiently inhibit *ZC3H12C* expression during cell manufacturing by, for example, RNP-delivery of CRISPR interference (CRISPRi) components, shRNAs, or small molecules inhibiting the ZC3H12C protein, which may be sufficient to generate a superior infusion product.

While our study establishes *ZC3H12C* as a target to improve T cell performance, several questions remain unanswered. First, the molecular mechanism by which *ZC3H12C* limits T cell therapy is not fully defined. *ZC3H12C* is a member of the Regnase family of RNA endonucleases, and may therefore function as a post-transcriptional regulator of inflammatory gene programs, analogously to the canonical *ZC3H12A* (Regnase-1)^48^. In this context, we speculate that *ZC3H12C* represents a Regnase-like checkpoint that is deployed in a state- and time-dependent manner to reshape the balance of T cell states^39,49–51^. However, the differential expression of *ZC3H12C* between acutely activated and exhausted T cell states suggests that Regnase-3 does not impact activation *per se* but could be preferentially associated with the establishment or maintenance of T cell exhaustion. Hence, Regnase-3 could represent a similar mode of action as other Regnase-family members^33,40–42^ albeit with a specific relevance to T cell exhaustion. Second, although improved efficacy in a metastatic as well as a transplanted solid tumor model is encouraging, additional studies across diverse solid tumor settings are required to validate *ZC3H12C* as a target with broad relevance. To this end, a study submitted for review in parallel to our work (Barbao et al.) identified *ZC3H12C* in an *in vivo* CRISPR screen and confirmed the efficacy of *ZC3H12C* engineered CAR T cells in diverse tumor models, supporting this gene as a platform-independent target to improve cell therapy. Third, while knockout improved antitumor activity in multiple models, long-term consequences of sustained enhancement of effector function including potential impacts on memory formation, exhaustion resistance versus terminal differentiation, and safety require careful evaluation.

Taken together, we identify a new role for *ZC3H12C* as a T cell dysfunction-specific remodeled gene that represents a platform-independent target to improve adoptive T cell therapy.

## Acknowledgements

We thank the members of the LIT FACS Core Facility for their excellent technical support. The research was supported by: Deutsche Forschungsgemeinschaft (DFG) (SCHM 3397/3-1 and SCHM 3397/3-2 (to C.S.), Projektnummer 324392634 – TRR 221 (to H.P. and C.S.), PO 1575/5-1 (to H.P.)), the German Cancer Aid (70114547 to H.P., 70115337 to C.S.), the Bavarian Cancer Research Center (BZKF) (to. M.P. and H.P.) and the European Union (project MICROBOTS, Grant No. 101124680 to H.P.); C.H.-L. and C.R.F.S. are supported by the “T-FITNESS” Grant funded by the European Union under Grant agreement Number 101070740.

## Author Contributions

Conceptualization: C.S., H.P.; Funding acquisition: C.S., H.P., L.G; Methodology: G.K., M.P., L.K., C.R.F.S., D.S., R.S., L.G., F.M., P.N., M.K.-S., M.K., T.V., L.B.; Formal analysis: G.K., M.P., C.S., C.H.-L., D.S., S.L., L.G., C.G.; Investigation: G.K., M.P., G.L-G., K.H, S.G., C.H.-L., L.K., C.R.F.S., R.S.; Project administration: C.S., C.H.-L., H.P.; Resources: C.S., L.G., H.P; Writing – original draft: C.S., G.K.; Writing – review & editing: all authors; Supervision: C.S., L.G., H. P., M.P., C.H.-L;

## Data availability statement

The experimental data and code generated in this study are available from Zenodo at https://zenodo.org/records/XXX. Processed scATAC-seq and scRNA-seq data from human TILs and in vitro–activated T cells from Riegel et al.^24^ were obtained from Zenodo (https://doi.org/10.5281/zenodo.7885745). Processed scRNA-seq data from Zhang et al.^28^ were obtained from the UCSC Cell Browser (https://cells.ucsc.edu/?ds=covid19-immuno). Processed bulk RNA-seq data from Yao et al.^30^ and Philip et al.^10^ were obtained from the Gene Expression Omnibus (GEO) under accession numbers GSE119942 and GSE89307, respectively. Processed scRNA-seq data from Haradhvala et al.^32^ were obtained from GEO (GSE197268) and from https://github.com/getzlab/Haradhvala_et_al_2022.

## Supplementary Figures and Legends

**Figure S1:**
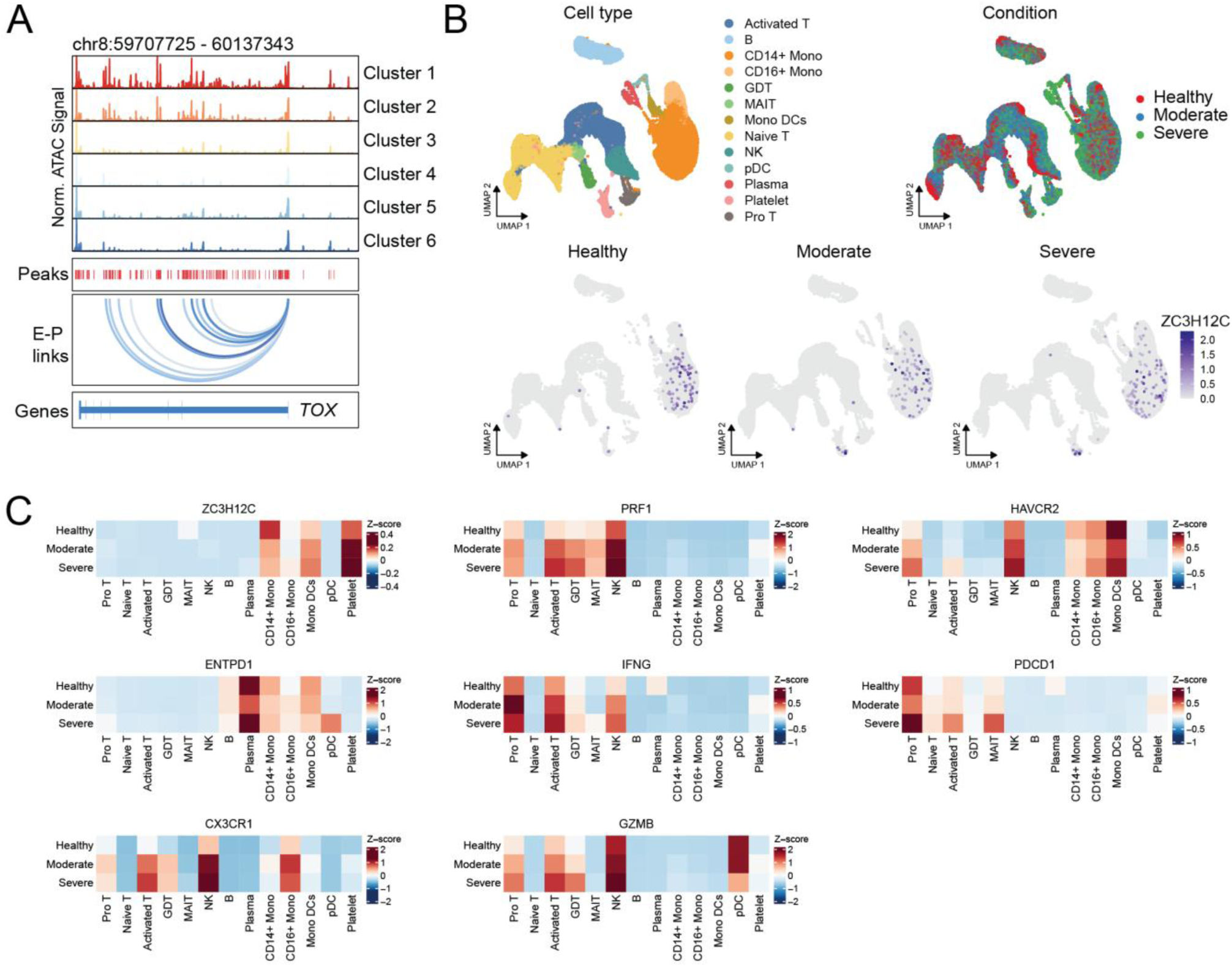
*ZC3H12C* is specifically induced in dysfunctional T cells. **(A)** Genome browser track visualization containing normalized chromatin accessibility (scATAC-seq) summarized for each cluster in Figure 1B, accessible regions (’peaks’), predicted regulation of gene promoters by distal enhancer peaks (‘E-P links’) with a value score indicating the correlation between accessibility of a peak and the expression of the corresponding gene, and the gene structure (’genes’) for the *TOX* locus. **(B)** Cluster annotation indicating PBMC cell subtypes and disease severity from the same experiment as reported in Figure 1G (upper panels). *ZC3H12C* expression in PBMCs stratified by disease severity, with healthy donors as controls (lower panels). **(C)** Z-score normalized expression of marker genes for each cell population in the blood, grouped into healthy donors, moderate, and severe COVID-19 patients.

**Figure S2:**
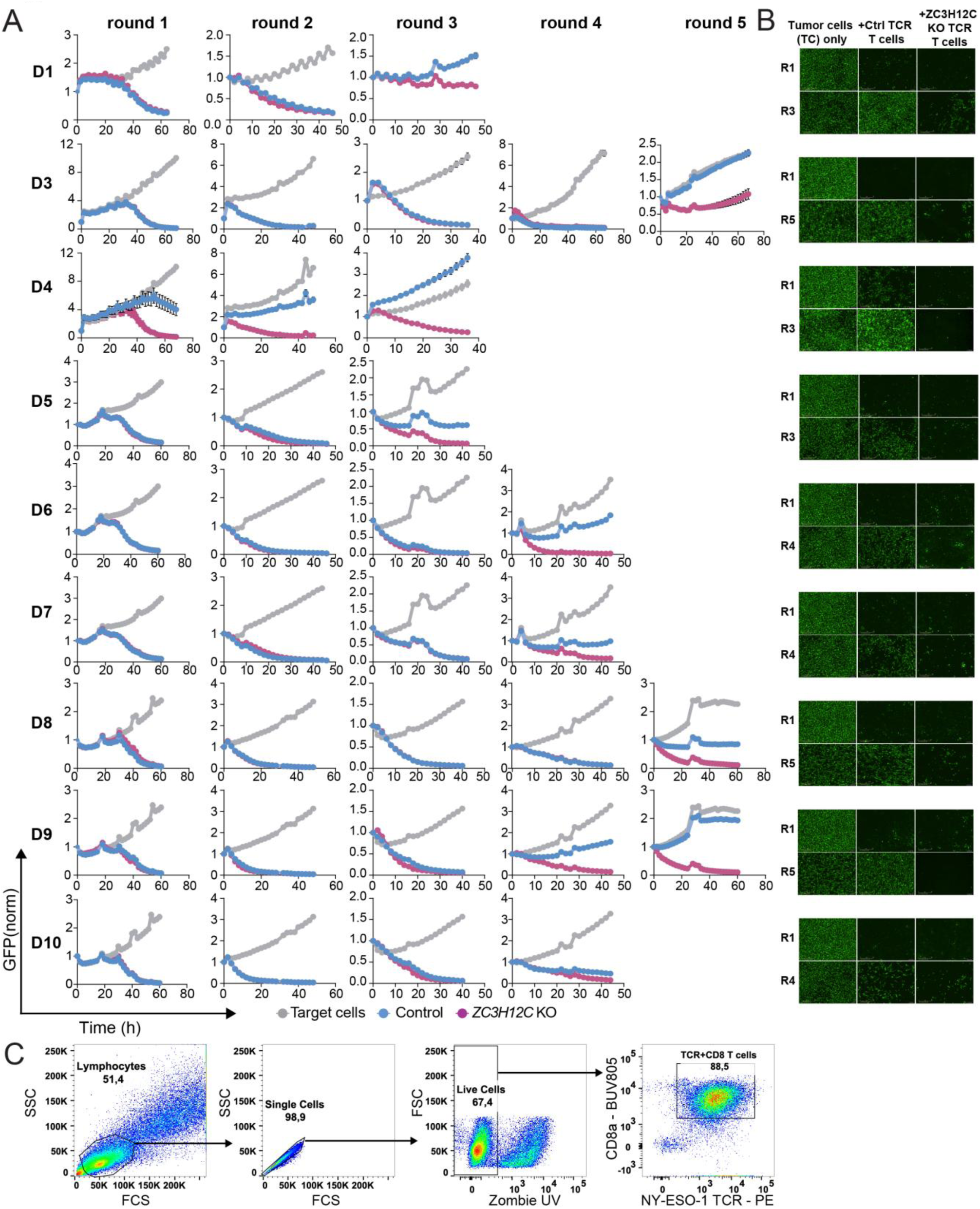
Impact of *ZC3H12C* deletion on cytotoxicity and effector molecule expression in NY-ESO-1 TCR T cells. **(A)** Real-time cytotoxicity kinetics over three to up to five consecutive rounds of NY-ESO-1 TCR T cell co-culture with SK-MEL-23 melanoma target tumor cells (E:T=1:1). T cell mediated cytotoxicity was quantified by loss of GFP signal from GFP^+^ SK-MEL-23 melanoma target cells. *ZC3H12C* KO TCR T cells were compared to Control *AAVS1* TCR T cells, tumor target cells only are shown as a further control. Ten biological replicates were performed, nine are shown here, donor 2 is shown in main Figure 2D. **(B)** Representative Incucyte images matching the cytotoxicity assays in (A) showing GFP^+^ target cells from round one and the round of co-culture determined as the endpoint when not enough Control *AAVS1* TCR T cells were left to continue a next round of co-culture. **(C)** Gating strategy of flow cytometry analysis in Figure 2F-G.

**Figure S3:**
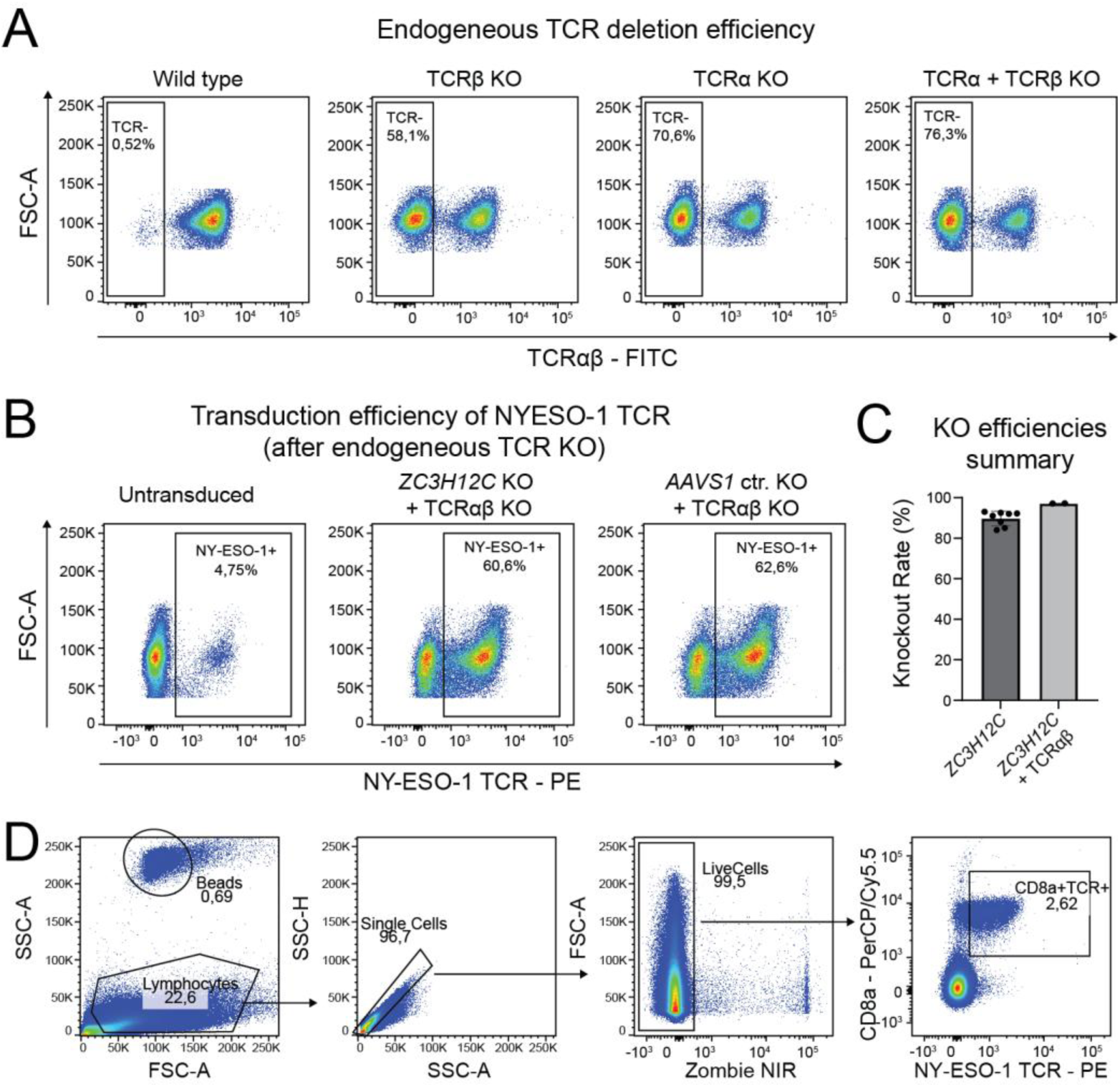
NY-ESO-1 TCR T cell CRISPR editing, TCR transduction, and flow cytometry gating strategy. **(A)** Establishment of simultaneous knockout of *TRAC* and *TRBC* in T cells prior to NY-ESO-1 TCR transduction. **(B)** Gating strategy of the flow cytometry analysis from Figure 3B-C. **(C)** Summary of *ZC3H12C* and combined *ZC3H12C* and endogenous TCR CRISPR knockout efficiencies as determined by the TIDE assay. **(D)** Gating strategy of the flow cytometry analysis in Figure 3G-F.

**Figure S4:**
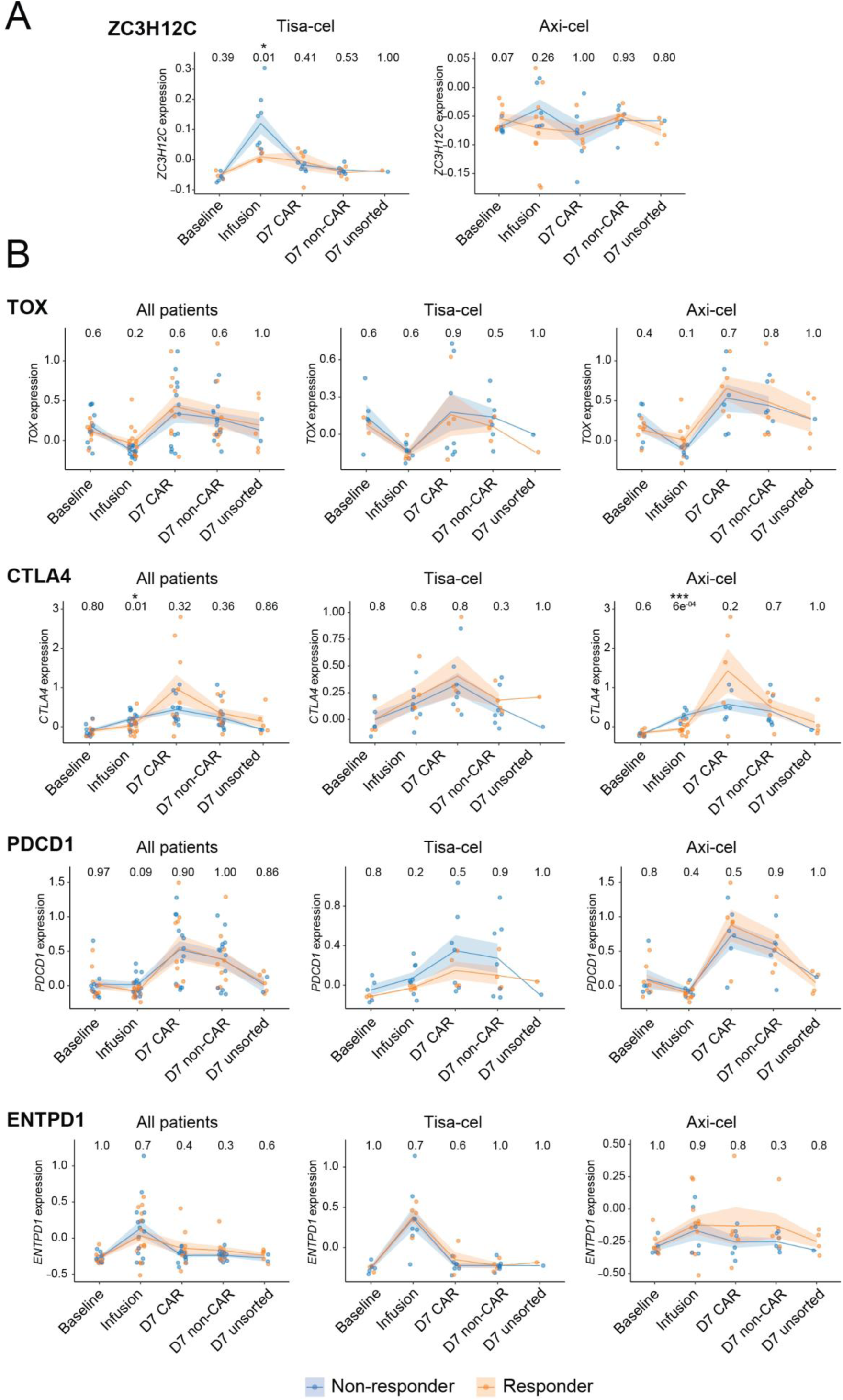
*ZC3H12C* is upregulated in pre-infusion CAR T cell products of non-responding patients. **(A)** *ZC3H12C* expression stratified by clinical response (responders versus non-responders) in T cells after isolation (Baseline), the CAR T cell infusion product (Infusion), CAR T cells isolated from patients seven days after infusion (D7 CAR), T cells without CAR expression day seven after infusion (D7 non-CAR), and unsorted T cells seven days after infusion (D7 unsorted). Data are grouped by CAR product type (Tisa-cel and Axi-cel). Each point represents the mean pseudobulk expression of a patient at each timepoint. **(B)** Expression of additional markers similar to (A). Statistical significance was assessed using a two-sided unpaired Wilcoxon rank-sum test. Numbers in the top row indicate p values.

## Methods

### Cell Culture

Pan T cells were isolated using Miltenyi MACS isolation (Miltenyi Biotec 130-096-535), from fresh or frozen buffy cones obtained from healthy donors at the Transfusion Medicine, University Hospital, Regensburg. Cells were cultured in RPMI 1640, GlutaMAX^TM^ (Gibco 61870036) supplemented with 10% heat-inactivated Fetal Calf Serum (FCS; Sigma Aldrich F7524) and 1% Penicillin Streptomycin (Gibco 15140122). CD8^+^ T cells were isolated using Miltenyi MACS isolation (Miltenyi Biotec 130-045-201) from fresh or frozen buffy cones obtained from healthy donors at the Transfusion Medicine, University Hospital, Regensburg. Cells were cultured in RPMI 1640, GlutaMAX^TM^ supplemented with 10% heat-inactivated FCS and 1% Penicillin Streptomycin. NY-ESO-1^+^ SK-MEL-23 melanoma cells stably expressing GFP were kindly provided by the laboratory of Luca Gattinoni. CD19^+^ NALM6 leukaemia cells stably expressing luciferase and HER2^+^ SKOV3 ovarian cancer cells stably expressing luciferase were kindly provided by the laboratory of Hendrik Poeck. All cell lines were cultured in complete RPMI 1640, GlutaMAX™ supplemented with 10% heat-inactivated FCS and 1% Penicillin Streptomycin. Lenti-X 293T cells were obtained from Takara Bio (632180) and maintained in DMEM supplemented with 10% heat-inactivated FCS and 1% Penicillin Streptomycin. Cells were passaged every third day using Trypsin EDTA. Ethics concerning experiments with human healthy donors were approved by the ethics committee of the University of Regensburg (reference numbers 19-1414-101 and 20-2040-101)

### Virus Production

#### HER2 CAR encoding lentivirus and NYESO-1 TCR encoding retrovirus production in HEK293T cells

HEK293T cells were seeded in 10 cm dishes at 3 million cells/dish in DMEM supplemented with 10% FCS and 1% Penicillin Streptomycin. Once the cells were 60-80% confluent, they were concomitantly transduced with helper plasmids (psPAX2 - Addgene 12260 and pMD2.G – Addgene 12259) and coding plasmid (Her2 CAR or NYESO-1 TCR coding plasmid) by using Lipofectamine 3000 (30 µl/dish). Virus-containing supernatants were collected 24 and 48 h after transduction. Virus was concentrated by ultracentrifugation for HER2 CAR-expressing lentivirus or 10 k MWCO Protein concentrator (Cytiva) for NY-ESO-1 TCR - expressing retrovirus. Virus titers were determined using Jurkat cells and 1.5-5 MOI of virus was used for transduction of primary T cells.

#### CD19 CAR encoding retrovirus production in PG-13 cells

PG-13 packaging cells were used to generate CD19-encoding retroviral particles. Briefly, PG-13 cells were transiently transduced with plasmids encoding the CD19 retroviral construct to initiate virus production. Following transduction, virus-containing supernatants were collected, and 500 µL of supernatant was used directly for subsequent transduction of target cells.

### Flow cytometry

Cells were stimulated with PMA/Ionomycin in the presence of protein transport inhibitors (BD Pharmingen™ Leukocyte Activation Cocktail, with BD GolgiPlug™ 550583 + BD GolgiStop*TM* 554724) for up to 6 h. Following stimulation, cells were washed in 1x PBS and stained with ZombieUV viability dye (BioLegend, 423108) to discriminate live and dead cells. Cells were then washed and stained with fluorophore-conjugated antibodies; BUV805-CD8a (BD Biosciences 742030), PE-TCRVβ13.1 (NYESO-1 TCR) (BioLegend 362410) and BB515-CD39 (BD Biosciences 565469) for 30 min on ice. After surface staining, cells were fixed and permeabilized using the eBioscience™ Foxp3/Transcription Factor Staining Buffer Set (eBioscience, 00-5523-00) according to the manufacturer’s instructions, followed by intracellular staining for APC-TOX (Miltenyi Biotec 130-118-474), PE-Cyanine7-EOMES (ThermoFisher Scientific 25-4877-42), BV650-Anti-T-bet (BD Biosciences 564142), APC/Fire*TM*750-Granzyme B (BioLegend 372210), BB700-IFNγ (BD Biosciences 566394), BUV395-IL-2 (Thermo Fisher Scientific 363-7029-42) and PE/Dazzle*TM*594 (BioLegend 502945). Samples were acquired on a BD FACSymphony A5 within 3 days of staining. Compensation controls were prepared using UltraComp eBeads™ Plus Compensation Beads (Thermo Fisher Scientific, 01-3333-41). Flow cytometry data were analysed using FlowJo v10. For mouse blood, samples were subjected to red blood cell lysis using 200 μl ACK lysis buffer (Gibco A1049201) followed by washing with 1x PBS. Cells were then stained with Zombie NIR™ fixable viability dye (BioLegend 423106) for 15 min at room temperature, washed with 1X FACS buffer, and subsequently incubated with fluorophore-conjugated antibodies: PerCP/Cyanine5.5 CD8 (BioLegend 344710), PE-TCRVβ13.1 (BioLegend 362410) for cell-surface marker staining. Data were acquired on a BD FACSymphony™ A5 flow cytometer. Absolute cell numbers were determined by addition of Precision Count Beads (BioLegend 424902) according to the manufacturer’s instructions.

### Electroporation

#### ZC3H12C KO in CAR T cells

CAR T cells were electroporated using the Lonza 4D-Nucleofector system and the P3 Primary Cell 4D-Nucleofector® X Kit S (Lonza V4XP-3032). For ribonucleoprotein (RNP) assembly, CRISPR RNAs (crRNAs; 200 µM) targeting *ZC3H12C* (GTATGGATACCGTTAATGTG and CAGTGCTCTCCACATAAAGG) or *AAVS1* (GACGCAAGGGAGACATCCGT and TGGAGAGGTGGCTAAAGCCA) were combined with trans-activating crRNA (tracrRNA; 200 µM; IDT 1072533) at a 1:1 molar ratio and annealed by heating to 95 °C for 5 min. Duplexed guide RNAs (gRNAs) were cooled at room temperature for 10–15 min, followed by addition of Cas9 nuclease (6 µg per gRNA; Thermo Fisher Scientific, A36499) and incubation at room temperature for 20 min to allow RNP complex formation. Pre-activated CAR T cells were resuspended in P3 nucleofection buffer at 2.5 x 10^6^ cells per 20 µl per well and combined with the pre-assembled RNPs in a 16-well Nucleocuvette^TM^. Electroporation was performed using program CA137, and cells were subsequently recovered in complete medium containing 25 IU/ml IL-7 (Miltenyi Biotec 130-095-367) and 50 IU/ml IL-15 (Miltenyi Biotec 130-095-760) for 3 days prior to downstream analyses.

#### ZC3H12C KO for TCR T cells

CD8^+^ T cells were activated using TransAct (1:100; Miltenyi Biotec, 130-111-160) in the presence of recombinant human IL-2 (100 IU/ml; Proleukin S, Novartis). At 40 h post activation, cells were electroporated using the Lonza 4D-Nucleofector system and the P3 Primary Cell 4D-Nucleofector® X Kit S. For RNP complex formation, crRNAs (200 µM) were combined with tracrRNA (200 µM) at a 1:1 molar ratio and annealed by heating to 95 °C for 5 min, followed by cooling at room temperature for 10-15 min. Cas9 nuclease (6 µg per gRNA) was then added and incubated at room temperature for 20 min to allow RNP assembly. Pre-activated CD8^+^ T cells were resuspended in P3 nucleofection buffer at 2.5 x 10^6^ cells per 20 µl per well and mixed with the corresponding RNPs targeting *ZC3H12C* or *AAVS1* in a 16-well Nucleocuvette^TM^. Electroporation was performed using program EH115, and cells were recovered in complete medium containing 100 IU/ml IL-2 for 6 h prior to viral transduction.

### TCR T cells manufacturing for *in vitro* experiments

CD8^+^ T cells were activated for 40 hours in 1:100 Transact + 100 IU/ml anti-human IL-2. Activated CD8^+^ T cells were then electroporated with Cas9/RNP complexes for either *ZC3H12C* or *AAVS1*. 6 hours post electroporation; these cells were then transduced with 3-5 MOI of NY-ESO-1 encoding retrovirus via spinoculation in a RetroNectin (Takara T100A) coated 24 well plate (Greiner 662160). Cells were allowed to rest for 3 days before transduction efficiency was determined via flow cytometry.

### CAR T cell manufacturing for *in vivo* experiments

#### HER2 CAR T cells

Pan T cells were activated for 72 hours in 1:100 Transact + 25 IU/ml anti-human IL-7 + 50 IU/ml anti-human IL-15. Activated Pan T cells were then transduced with 1.5 MOI of HER2 CAR encoding lentivirus via spinoculation in a RetroNectin-coated 24 well plate. CAR expression was assessed by flow cytometry 24 hours post-transduction. Following transduction, cells were electroporated with Cas9 ribonucleoprotein (RNP) complexes targeting *ZC3H12C* or with Cas9-only control RNPs. Cells were rested for 4 days, after which knockout efficiency was evaluated by Sanger sequencing.

#### CD19 CAR T cells

Pan T cells were activated for 48 hours in 1:100 Transact + 25 IU/ml anti-human IL-7 + 50 IU/ml anti-human IL-15. Activated Pan T cells were then transduced with 500 µl of viral supernatant containing CD19 CAR encoding retrovirus via spinoculation in a RetroNectin-coated 24 well plate. The following day, transduced T cells were harvested and a following second transduction was performed with the previously transduced T cells, as mentioned above. CAR expression was assessed by flow cytometry after transduction on the same day. Following transduction, cells were electroporated with Cas9 ribonucleoprotein (RNP) complexes targeting *ZC3H12C* or with Cas9-only control RNPs. Cells were rested for 4 days, after which knockout efficiency was quantified by Sanger sequencing.

### TCR T cell manufacturing for *in vivo* experiments

To avoid a strong GvHD in mice, TCR T cells were produced with a knockout for endogenous TCR as described previously^52^. CD8*+* T cells were activated for 40 hours in 1:100 Transact + 100 IU/ml anti-human IL-2. Activated CD8*+* T cells were then electroporated with Cas9/RNP complexes for single gRNA for TCRα (5′-AGAGTCTCTCAGCTGGTACA-3′) and TCRβ (5′-GGAGAATGACGAGTGGACCC-3′) concomitantly with either *ZC3H12C* or *AAVS1*. 24 hours post electroporation, these cells were then transduced with 3-5 MOI of NY-ESO-1 encoding retrovirus via spinoculation in a RetroNectin-coated 24 well plate. Cells were allowed to rest for 3 days before knockout efficiency and transduction efficiency was determined via flow cytometry using PerCP/Cyanine5.5 CD8 (BioLegend 344710), PE-TCRVβ13.1 (NYESO-1 TCR) (BioLegend 362410) and FITC-TCR α/β (BioLegend 306706) antibodies. Further enrichment was carried out via flow cytometry-based cell sorting to ensure 70-100% TCR*+* T cells.

### Sanger Sequencing and TIDE Assay

0.5-1 million cells with or without CRISPR knockout for *ZC3H12C* were collected and DNA was isolated from these cells using the DNeasy Blood & Tissue Kit (Qiagen 69506). PCR amplification of the KO site was carried out by using 5’-CCATTGAGACAGAAATCTAGCCA-3’ and 5’-GTATTCTCGCACCACATCAGG-3’. Amplified product was then cleaned up using PCR Cleanup kit (Qiagen 28104) and sent for Sanger sequencing using the same primers. DNA sequences were then analysed using TIDE^53^ (https://tide.nki.nl/) to determine indels around the cut site.

### Killing and Restimulation assay

To assess antigen-dependent expansion and serial cytotoxic function of NY-ESO-1 TCR-engineered T cells, a serial re-stimulation co-culture assay was performed using GFP-expressing SK-MEL-23 melanoma target cells. NY-ESO-1^+^ TCR T cells were co-cultured with GFP^+^ SK-MEL-23 cells at an effector-to-target (E:T) ratio of 1:1 in complete RPMI medium (without IL-2) in a 24 well flat bottom clear plate. Co-cultures were maintained for 48 hours per round and monitored by live-cell imaging (Incucyte, Sartorius) to quantify target cell killing based on the loss of GFP signal. At the end of each round, T cells were harvested, counted, and plated onto freshly seeded GFP^+^ SK-MEL-23 target cells, maintaining the E:T ratio of 1:1, to initiate the next round of antigen re-challenge. Serial re-stimulation was repeated for up to five consecutive rounds. T cells were collected after round 3 for flow cytometry analysis.

### Mouse Experiment

All mice were housed under SPF conditions in individually ventilated cages at the Animal Center of University Hospital Regensburg. All experiments were approved by the Animal Ethics Committee of University Hospital Regensburg and the Animal Ethics Committee of the Government of Lower Franconia (Animal Protocol Numbers: RUF-55.2.2-2532-2-1563, RUF-55.2-2532-2-1838, and RUF-55.2.2-2532-2-1610) and performed in accordance with institutional and national guidelines.

#### Xenograft CD19 CAR T cell model

Immunodeficient female 10- to 12-week-old NOD-*Prkdc^scid^IL2rg^Tm^*^1^/Rj (Janvier Labs, France) were injected i.v. into the tail vein with 0.5 x 10^6^ CD19^+^ Nalm-6/EGFP-Luc 7 days prior to CAR T cell injection (day -7). On day 0, mice were randomized into equal groups based on bioluminescence measurements. 1 x 10^6^ anti-CD19-CAR T cells with or without the knockout for *ZC3H12C*, were injected i.v. into the tail vein. Treatment with 0.5 x 10^6^ untransduced Pan T cells served as control. Mice were scored daily. Tumor progression, systemic seeding, and response to CAR T therapy was confirmed regularly by bioluminescence imaging using a Xenogen IVIS imaging system (PerkinElmer, United States) after i.p. injection of 200 µl IVISbrite D-Luciferin (15 mg/ml, Revvity; United States), and the intensity of the signal was measured as average radiance (photon/second/cm^2^/steradian [p/s/cm^2^/sr]) and total flux (photons/second [photons/s]). Mice were euthanized utilizing humane endpoints or 100 days after (CAR) T cell injection.

#### Xenograft HER2 CAR T cell model

Immunodeficient female 10- to 12-week-old NOD-*Prkdc^scid^IL2rg^Tm^*^1^/Rj (Janvier Labs, France) were injected i.v. into the tail vein with 1.5 x 10^6^ HER2^+^ SKOV3/EGFP-Luc 7 days prior to (CAR) T cell injection (day -7). On day 0, mice were randomized into equal groups based on bioluminescence measurements. To evaluate the effect of *ZC3H12C* KO, 1 x 10^6^ anti-HER2-CAR T cells with or without the knockout for *ZC3H12C*, were injected i.v. into the tail vein. Treatment with 0.5 x 10^6^ untransduced Pan T cells served as control. Mice were scored daily. Tumor progression, systemic seeding, and response to CAR T therapy was confirmed regularly by bioluminescence imaging using a Xenogen IVIS imaging system (PerkinElmer, United States) after i.p. injection of 200µl IVISbrite D-Luciferin (15 mg/ml, Revvity; United States), and the intensity of the signal was measured as average radiance (photon/second/cm^2^/steradian [p/s/cm^2^/sr]) and total flux (photons/second [photons/s]). Mice were euthanized utilizing humane endpoints or 100 days after (CAR) T cell injection.

#### NY-ESO-1 TCR T cell model

1×10^6^ NYESO-1+ SK-MEL-23 melanoma cells in PBS were injected subcutaneously into the shaved flank of 10- to 12-week-old female NOD.Cg-*Prkdc^scid^ Il2rg^tm1Sug^* Tg(CMV-IL2/IL15)1-1Jic/JicTac (Taconic Biosciences) 27 days prior to (TCR) T cell injection (day - 27). On day 0, mice were allocated to groups based on tumor size and injected with 2 x 10^6^ NY-ESO-1-TCR T cells in PBS intravenously. Tumor width (w) and height (h) were measured using calipers and tumor area (T) was calculated as T = w × h. Peripheral blood samples were taken from the lateral tail vein.

### Analysis of *ZC3H12C* chromatin accessibility and expression across public datasets

Analyses of the public datasets were performed in R (v4.3.4) using the GEOquery (v2.74.0), Seurat^54^ (v5.3.0), SeuratExtend^55^ (v1.2.7), and ArchR^56^ (v.1.0.3) packages, and in Python (v3.12.11) using Scanpy^57^ (v1.9.3), and ADPbulk (v0.1.4).

#### Gene prioritization analysis in dysfunctional TILs

Ranking of genes based on exhaustion-specific chromatin accessibility (super enhancer score, ‘SE score’, Figure 1A), was derived from Riegel et al. In brief, genes were ranked and plotted based on the number of T_EX_-cluster-specific enhancer-promoter interactions.

#### Integrated scATAC-seq and scRNA-seq from human TILs and in vitro activated T cells

Processed sequencing data from Riegel et al.^24^ was obtained from ZENODO (https://doi.org/10.5281/zenodo.7885745). For UMAP projection and marker gene visualization, the ‘Core’ analysis object was used (ArchRobject_TIL_core.zip), and marker genes based on chromatin-based gene activity scores, or gene expression based on integrated scRNA-seq datasets were plotted using ArchR. Genome browser tracks were also created using ArchR. For the comparison of *ZC3H12C* activity *in vitro* activated versus *in vivo* exhausted TILs, the analysis object containing also *in vitro* activated cells was used (‘ArchRobject_TIL_all.zip’). Here, all T_EX_ clusters were merged and compared against in vitro activated CD8^+^ T cell cluster form healthy donor PBMCs.

#### scRNA-seq analysis of COVID-19 patient PBMCs

scRNA-seq data from Zhang et al.^28^ were analyzed to assess *ZC3H12C* expression in peripheral blood during acute SARS-CoV-2 infection. Processed and normalized gene expression data with cell-level metadata and UMAP coordinates were obtained from the UCSC Cell Browser. Hematopoietic stem cells were excluded due to low abundance. Analyses focused on healthy donors (n=5; 22,681 cells), moderate COVID-19 cases (n=7; 37,880 cells), and severe COVID-19 cases (n=4; 24,624 cells). To compare expression across conditions, mean gene expression was computed per cell type and condition, then gene-level Z-scores were calculated across conditions.

#### Bulk RNA-seq analysis of acute vs chronic LCMV infection

Bulk RNA-seq data from Yao et al.^30^ were analyzed to examine *Zc3h12c* expression in mouse P14 CD8^+^ T cells during acute versus chronic infection. The dataset included three replicates per condition from days 4.5 and 7 after infection with either the acute Armstrong or chronic clone 13 LCMV strain. Preprocessed TMM-normalized expression data were retrieved from GEO (GSE119942), and *Zc3h12c* expression was compared across time points within each infection group.

#### Bulk RNA-seq analysis of acute vs tumor-infiltrating T cells

Bulk RNA-seq data from Philip et al.^10^ were analyzed to assess *ZC3H12C* expression in mouse CD8^+^ T cells across differentiation states. The acute infection setting included naïve, effector (days 5 and 7), and memory T cells, while the tumorigenesis setting TILs collected at days 5, 7, 14, 21, 28, 35, and 60. Each condition included three replicates. Normalized count data were obtained from GEO (GSE89307), and *Zc3h12c* expression was compared across time points within each setting.

#### scRNA-seq analysis of CAR T cell therapy in refractory B-cell lymphoma

scRNA-seq data from Haradhvala et al.^32^ were analyzed to examine the association between *ZC3H12C* gene expression and clinical response in 32 patients with refractory large B-cell lymphoma treated with CAR T cell therapy (non-responder n=15; responder n=16). Samples were collected at baseline (PBMCs), from the CAR T infusion product, and at day 7 post-infusion, with day 7 samples sorted into CAR-positive, CAR-negative, or unsorted fractions. Processed and normalized gene expression data with cell-level metadata and UMAP coordinates were obtained from the published dataset. Analyses were restricted to CD4^+^ and CD8^+^ T cells, yielding 235,683 cells in total (36,069 baseline; 94,544 infusion; 34,667 day 7 CAR-positive; 47,377 day 7 CAR-negative; 23,026 day 7 unsorted). Pseudobulk expression profiles were generated per patient and treatment stage by averaging normalized gene-expression values across T cells and used to visualize *ZC3H12C* expression over the treatment course.

## Statistical Analysis

Statistical analyses were performed using GraphPad Prism. For comparisons between control and KO groups, statistical significance was determined using a paired two-tailed Student’s t-test. Survival curves were analyzed using the log-rank (Mantel–Cox) test. P < 0.05 considered statistically significant.

